# Sequencing and functional analysis of *Sphingobium yanoikuyae* A-TP genome reveals genes for utilization of limonene, α-pinene, and citronellol

**DOI:** 10.1101/2025.04.30.651573

**Authors:** Nemanja Jankovic, Giovanni Marques de Castro, André Luiz Quintanilha Torres, Rafael Chelala Moreira, Marcela Uliano-Silva, Cinara Souza da Conceição, Vitor Lima Coelho, Juliana Alves Americo, Juliano Lemos Bicas, Mauro de Freitas Rebelo

**Affiliations:** Carlos Chagas Filho Institute for Biophysics, Federal University of Rio de Janeiro, Brazil; Department of Innate Immunity, UMass Chan Medical School, USA; Department of Entomology, University of California, Riverside, USA; Department of Food Science and Nutrition, School of Food Engineering, State University of Campinas, Brazil; Wellcome Sanger Institute, Cambridge, UK; SENAI CETIQT, Rio de Janeiro, Brazil; Bio Bureau Biotechnology, Rio de Janeiro, Brazil

**Keywords:** Monoterpenes, metabolism, gene discovery, functional genomics, biotechnology, limonene 8-hydratase

## Abstract

*Sphingobium yanoikuyae* A-TP is a bacterial strain capable of utilizing and biotransforming commercially relevant monoterpenes. Genome mining and similarity-based analysis identified two gene clusters (24 and 30 genes) likely responsible for limonene, α-pinene, and citronellol metabolism. Notably, genes associated with α-pinene metabolism — an oxide lyase and an aldehyde dehydrogenase — were clustered with those for citronellol utilization, despite limonene’s closer structural similarity to α-pinene. Growth experiments confirmed citronellol as a carbon source, but linalool, geraniol and β-myrcene were not substrates for this strain. To address the unresolved identity of limonene 8-hydratase, the enzyme catalyzing limonene conversion to α-terpineol, we performed RNA-seq analysis on strains with contrasting phenotypes which identified highly upregulated candidate genes. Phylogenetic analysis supports the classification of this strain as a *Sphingobium yanoikuyae* species. This study advances the understanding of monoterpene catabolism in *Sphingomonadaceae* and provides a foundation for biotechnological production of monoterpenes and their derivatives.

**Impact statement:** Monoterpenes, such as limonene and *α*-pinene, are often used as starting material to produce derivatives that are of interest to the aroma and fragrance industry. In this work we uncover the genes harbored in the *Sphingobium yanoikuaye* A-TP genome that are putatively involved in the metabolism of limonene, citronellol and *α*-pinene. We also propose several candidates for the limonene 8-hydratase, an enzyme that converts limonene to *α*-trepineol. The discovery of these genes can be explored for biotechnological production of industrially important compounds.

**Data summary:** All genomic and transcriptomic data from Illumina sequencing are available at the NCBI GenBank archive under the BioProject accession number PRJNA1196986. The Supplementary Material has been uploaded to Microbiology Society figshare: 10.6084/m9.figshare.28184306

**Author Notes:** The authors confirm all supporting data, code and protocols have been provided within the article or through supplementary data files. Seven supplementary figures, three supplementary tables, and four supplementary files are available with the online version of this article.

## Introduction

Volatile compounds, such as monoterpenes (C10) and sesquiterpenes (C15), are among the most valuable fragrance compounds due to their distinct scent (1). Terpenes are specialized metabolites that play essential roles in ecological interactions of plants, including defense against herbivores and communication with other plants (2) However, their chemical synthesis might be difficult and costly, while direct extraction from the natural sources is typically low yielding (1).

An alternative approach is microbial production of terpenes, either through *de novo* biosynthesis in engineered hosts using inexpensive carbon sources, or by whole-cell biotransformation when structurally related precursors are readily available. Deciphering metabolic pathways from microorganisms that have evolved to break down monoterpenes and use them as a source of energy can help expand the current synthetic biology tools for the production of natural fragrance compounds (1).

The metabolic pathways have been revealed for several bacterial species with the ability to break down cyclic (limonene, α-pinene) and acyclic (citronellol, β-myrcene) monoterpenes, such as *Rhodococcus erythropolis* (3), *Pseudomonas rhodesiae* (4), *Pseudomonas aeruginosa* PAO1 (5), and *Pseudomonas* sp. M1 (6). For instance, *P. aeruginosa* PAO1 harbors the well-characterized acyclic terpene utilization (*atu*) gene cluster, which converts citronellol into acetyl-CoA (5) (figure S1), while a gene cluster was described in *Pseudomonas* sp. M1 strain that allows β-myrcene utilization (6).

*Sphingobium* sp. has also been identified as an efficient monoterpene degrader that can grow on limonene, and *α*- and β-pinene (figure S2 and S3) (7). This strain can also convert limonene into α-terpineol, a commercially important aroma and fragrance compound, via a separate biotransformation pathway (figure S2) (8,9). However, the gene encoding the enzyme responsible for this conversion, limonene 8-hydratase, remains unknown. In addition to this, the genetic basis for utilization of limonene and pinene by this strain is also unresolved. We conducted a genomic and transcriptomic study of *Sphingobium* sp. to identify monoterpene degradation genes, to uncover candidate genes encoding limonene 8-hydratase and to evaluate the strain’s ability to utilize selected acyclic monoterpenes. We also performed a phylogenetic analysis to resolve the taxonomic classification of this strain.

## Material and methods

### DNA extraction and genome sequencing of *Sphingobium* sp

A single colony of *Sphingobium* sp. was inoculated in 25 mL of tryptic soy broth and incubated for 17 h at 30 °C and 200 rpm. Genomic DNA was isolated with the DNeasy Blood & Tissue DNA kit (QIAGEN) according to the supplier’s instructions and stored in 100 µL of TE buffer (10 mM Tris 1 mM EDTA, pH 8.0). The purity and concentration of the isolated DNA were evaluated on Nanodrop and with the Qubit dsDNA BR Assay kit (Thermo Scientific), respectively, while the DNA integrity was verified by agarose gel electrophoresis. The libraries were prepared with 500 bp inserts for 250 bp paired-end reads. Genome sequencing was conducted on Illumina HiSeq 2000 platform with an approximated yield of 2 Gb. This corresponds to approximately 400X coverage, assuming a genome size of approximately 5 Mb.

### Genome assembly and functional annotation

After sequencing, adapters were removed via Trimmomatic v.0.32 (10). The quality of the reads was analyzed with the Fastqc v.0.11.5 (11), and the genome was assembled with Spades v.3.10.1 (12). The corresponding genes and proteins were predicted and functionally annotated with RAST (13). The completeness of genome assembly and protein prediction was evaluated with the BUSCO v.3 algorithm (14). Annotation of *Sphingobum* sp. genes and proteins was conducted using KAAS (15) and QuickGO web servers (16), and Blast2GO v.5.2 (17). Conserved domains, transmembrane domains and signal sequences were predicted with InterProScan v.5.33 (18), DeepTMHMM v.1.0.8 (19) and SignalP 5.0 (20), respectively.

The functional annotation of the predicted *Sphingobium* sp. protein dataset was additionally inferred by comparison with the proteins from monoterpene-metabolizing microorganisms: *Rhodococcus erythropolis* DCL14 (AJ272366.1), *Pseudomonas rhodesiae* CIP 107491 (MF946559.1), *Pseudomonas aeruginosa* PAO1 (AE004091.2), *Pseudomonas* sp. M1 (CP094343.1) and *Grosmannia clavigera* kw1407 (GL629729.1, NW_014040759.1), using BLAST algorithm (21). The genomic data has been deposited to the NCBI database under accession number: JBJYKA000000000.

### Phylogenomic analysis

A comprehensive taxonomic analysis based on whole-genome data was conducted by submitting the strain’s genome sequence to the Type (Strain) Genome Server (TYGS) (22). After determining closely related type strains, all pairwise comparisons among the genomes were conducted using Genome Blast Distance Phylogeny (GBDP) and accurate intergenomic distances inferred under the algorithm ‘trimming’ and distance formula d_5_ (22). The results were retrieved from TYGS on 30 October 2024.

### Growth on acyclic monoterpenes

The pre-cultures were prepared by inoculating a loopful (10 µL) of *Sphingobium* sp. cells into 10 mL of YM medium (BD Difco), supplemented with 2 % (v/v) of either citronellol (Merck, 98 %), geraniol (Aldrich, 98 %), β-myrcene (Merck, 85 %) or linalool (Fluka, 97 %), in three biological replicates. Following 48 h incubation at 30 °C and 150 rpm, 100 μL of the pre-cultures was used to inoculate 10 mL of Pseudomonas basal medium without carbon source (8), supplemented by 1 % (v/v) of the same monoterpenes. The cultures were incubated under the same conditions as precultures. An aliquot (100 μL) of each preculture and culture was transferred onto YM agar plates and homogenized with the Drigalski loop. The colonies were enumerated following incubation of Petri dishes (72 h, 30 °C).

### Gene expression by two *Sphingobium* sp. phenotypes in response to limonene biotransformation

The transcriptional profiles of the *Sphingobium* sp. phenotype with retained (Lim^+^) and lost (Lim^-^) ability to convert limonene to α-terpineol (see Results) were compared following limonene biotransformation procedure as described by Bicas et al. (2010) (8). Briefly, a single colony of *Sphingobium* sp. was inoculated, in three biological replicates for each phenotype (Lim^+^ or Lim^-^) in 250 mL of Pseudomonas basal medium containing 4 % (m/v) glucose as a sole carbon source (8). Liquid cultures were incubated at 30 °C until OD_600_ of 1.0, harvested by centrifugation (2600 g, 4 °C, 10 min) and resuspended in 20 mM phosphate buffer (pH 7.5) for approximately 7-fold concentrated suspension. The suspensions (25 mL) were transferred to Erlenmeyer flasks containing equal volume of *n*-hexadecane, forming a two-phase system. Limonene was added to a final concentration of 40 g L^-1^ of *n*-hexadecane and incubated at 30 °C and 200 rpm for 24 h (8).

#### Total RNA extraction and library construction

Aqueous phase aliquots (200 µL) of Lim^+^ and Lim^-^ suspensions from the previous step, were harvested by centrifugation (12000 g, 4 °C for 5 min) and the cells were immediately frozen in liquid nitrogen. Total RNA was extracted with TRIzol reagent (Invitrogen, USA), purified using a magnetic bead kit (Agencourt RNAClean XP, Beckman-Coulter) according to the manufacturer’s instructions, and eluted in 20 µl of RNase-free water. The purity and concentration of RNA was evaluated with Nanodrop and the Qubit RNA High Sensitivity kit (Thermo Scientific), respectively. The integrity of the isolated RNA was confirmed visually by denaturing gel electrophoresis (23). Libraries were constructed using the Stranded Total RNA Prep with Ribozero kit (Illumina), quantified using the Qubit DNA High Sensitivity kit (Thermo Scientific) and fragment sizes visualized by Tapestation (Agilent Technologies) with the D1000 ScreenTape System kit.

#### RNA sequencing and transcriptome assembly

RNA sequencing was performed on the Illumina NextSeq 550 platform, for 150 bp reads (2 x 75 bp). The quality control of the raw reads was performed with Fastp v.0.21.0 (24). Indexing and alignment of reads to the *Sphingobium* sp. genome (JBJYKA000000000) was performed with STAR v.2.7.9a (25) and the transcripts were assembled with the StringTie package v.2.1.7 (26) The transcripts were quantified using Salmon v.1.6.0 (27) and had their function predicted using the same set of annotation tools as used in the DNA functional analysis.

#### Differential expression analysis

Data normalization (Trimmed Mean of M) and Fisher’s exact test were performed with the EdgeR package v.3.13 (28) on the final transcriptome assembly (NCBI accession number: GLCA00000000). Transcripts were considered differentially expressed for the False Discovery Rate (FDR) and log fold change conditions: FDR <10^-9^ and |log_2_FC| > 4.

### Identification of the volatile metabolites

Volatile metabolites produced during trials with *Sphingobium* sp. cells were detected by gas chromatography-mass spectroscopy (GCMS-QP2020, Shimadzu, Japan). For the cells grown on acyclic monoterpenes, 1 mL culture aliquots were taken every 24 h for 5 days. Following centrifugation (10000 g, 4 °C, 5 min), the supernatants were extracted with ethyl acetate (1:1 vol ratio) and dried over Na_2_SO_4_. In case of limonene biotransformation by Lim^+^ and Lim^-^ cells, 1 mL samples were withdrawn from the *n*-hexadecane layer following 24 h incubation. The samples were injected in split mode (1 μL) and separated in a fused silica Agilent DB5 capillary column (30m, 0.25 mm id, 0.25 m). The mass spectrometer was operated at an interface temperature of 200 °C, electron ionization at 70 eV, and a mass scan range of 40-500 m/z. Helium was used as carrier gas at a flow rate of 1.0 mL min^-1^. The data obtained were processed using the GCMS Solution Postrun workstation v.2.41 (Shimadzu, Japan). The compounds were positively identified by comparing isotope mass spectra with the NIST 2011 library considering > 90% similarity.

## Results and Discussion

### Comprehensive genome assembly of *Sphingobium* sp. unveils regulatory networks and biosynthetic gene clusters for specialized metabolites

The assembled *Sphingobium* sp. genome is 5.6 Mb long and encodes 5,583 proteins (Table 1 and Supplementary file S1). For comparison, the reference genomes of other *Sphingobium* species’ are 3.1 to 6.6 Mb long and contain 2,968 to 6,053 protein-coding genes ([NCBI *Sphingobium* genomes](https://www.ncbi.nlm.nih.gov/datasets/genome/?taxon=165695&reference_only=true)). We assigned a function to 4,484 (80.3%) *Sphingobium* sp. proteins by at least one of the annotation methods (Table 1, figure S4). The BUSCO analysis revealed the presence of 96 % of the orthologous genes conserved in bacteria (Table 1) indicating a satisfactory genome assembly and protein prediction (14). Additional information regarding *Sphingobium* sp. genome assembly can be found in table S1.

**Table 1.** The list of key characteristics of the genomic sequence pertaining to the *Sphingobium* sp. strain. BUSCO score abbreviations: C - complete, S - single, D - duplicated, F - fragmented, M - missing, n - total number of orthologous clusters used in the BUSCO assessment.

| Feature | Value |
| --- | --- |
| Genome assembly size | 5,669,148 bp |
| Contigs | 310 |
| G+C content | 63.9% |
| tRNA genes | 56 |
| rRNA genes | 127 |
| Protein coding genes | 5,583 |
| Annotated proteins | 4,484 (80.3% total) |
| Hypothetical proteins | 1,099 (19.7% total) |
| BUSCO score | C: 96.0% (S: 95.5%, D: 0.5%), F: 0.5%,<br>M: 3.5%, n: 221 |

Attempts to isolate plasmid from this strain were unsuccessful (data not shown), suggesting that the monoterpene catabolic genes are chromosomally encoded in this strain as previously reported (29), even though many members of the *Sphingomonadaceae* family carry xenobiotic degradative plasmids (30).

Phylogenomic analysis using the TYGS web server (figure S5) confirmed that *Sphingobium* sp. is closely related to *Sphingobium yanoikuyae* ATCC 51230^T^, with strong genomic similarity (bootstrap = 100). A complementary 16S rRNA maximum-likelihood analysis further supported this classification, clustering the strain with 15 other *S. yanoikuya*e isolates (figure S6). Based on these findings, we designate the strain in the present study as *S. yanoikuyae* A-TP, highlighting its α-terpineol-producing ability.

### Identification of two genomic clusters involved in metabolism of monoterpenes

Genome analysis of *S. yanoikuyae* A-TP revealed two distinct monoterpene degradation clusters (Table 2). The first, a 25-kb cluster (*peg*.*3034–peg*.*3057*) (figure 1) contains genes involved in limonene metabolism, including limonene monooxygenase (*peg*.*3051*), limonene 1,2-diol dehydrogenase (*peg*.*3042*), and Baeyer-Villiger monooxygenase (*peg*.*3039*). Notably, a putative xylonolactonase (*peg*.*3046*) was also identified, suggesting a possible role in lactone ring-opening, a step previously considered spontaneous (31). This cluster is flanked by a mobile genetic element (*peg*.*3058*) and conjugal transfer genes (*peg*.*3028–peg*.*3030*), indicating that this cluster was potentially obtained through horizontal gene transfer.

**Table 2.** Putative genes, their products, and assigned functions involved in the catabolism of limonene, *α*-pinene, and citronellol by *S. yanoikuyae* A-TP strain. The table presents gene products as annotated by the RAST server, as well as BLASTp metrics (e-value and percent identity) obtained by comparing proteins from monoterpene-degrading organisms against the *S. yanoikuyae* A-TP protein set. Symbol “-” indicates no similarity to the query sequence. In the “Pathway” column, “Liu” and “Atu” refer to proteins associated with the leucine/isovalerate and citronellol pathways, respectively.

| Locus tag | Product (RAST) | BLASTp analysis of proteins involved in metabolism of monoterpenes |  |  |  |  |  |  |
| --- | --- | --- | --- | --- | --- | --- | --- | --- |
|  |  | Pathway | Query organism | Query protein | Acc. num. | Putative function in monoterpene metabolism | e-value | Identity (%) |
| *peg.3039* | Cyclohexanone monooxygenase | Limonene | *G. clavigera* kw1407 | Cyclohexanone monooxygenase (CMQ_6956) | EFX06635.1 | 1-Hydroxy-2-oxo limonene 1,2-monooxygenase | 7e-105 | 34.19 |
| *peg.3042* | 3-alpha-(or 20-beta)-hydroxysteroid dehydrogenase | Limonene | *G. clavigera* kw1407 | Short chain dehydrogenase reductase (CMQ_238) | EFW99920.1 | DCPIP-dependent limonene 1,2-diol dehydrogenase | 1e-29 | 32.83 |
| *peg.3051* | Monooxygenase | Limonene | *R. erythropolis* DCL14 | Limonene monooxygenase (LimB) | CAC20855.1 | Limonene monooxygenase | 3e-23 | 28.07 |
| *peg.3120* | Coniferyl aldehyde dehydrogenase | α-Pinene | *P. rhodesiae* CIP 107491 | Coniferyl aldehyde dehydrogenase | AUL81537.1 | Isonovalal dehydrogenase | 0.00 | 54.49 |
| *peg.3140* | Ester cyclase | α-Pinene | *P. rhodesiae* CIP 107491 | α-Pinene oxyde lyase | AUL81538.1 | α-Pinene oxide lyase | 1e-07 | 28.83 |
| *peg.3121* | CoA ester lyase | Atu | *P. aeruginosa* PAO1 (AtuA) | 3-Hydroxy-3-iso-hexenylglutarylCoA:acetate-lyase (AtuA) | AAG06274.1 | 3-Hydroxy-3-iso-hexenylglutarylCoA:acetate-lyase | - | - |
| *peg.3128* | Short chain acyl-CoA dehydrogenase | Atu | *P. aeruginosa* PAO1 (AtuD) | Citronellyl-CoA desaturase (AtuD) | AAG06277.1 | Citronellyl-CoA-dehydrogenase | 1e-58 | 30.53 |
| *peg.3129* | Citronellol and citronellal dehydrogenase | Atu | *P. aeruginosa* PAO1 (AtuB) | Citronellol/citronellal dehydrogenase (AtuB) | AAG06275.1 | Citronellol dehydrogenase | 4e-45 | 41.63 |
| *peg.3136* | Long-chain-fatty-acid--CoA ligase | Atu | *P. aeruginosa* PAO1 | Very long chain acyl-CoA synthetase (AtuH) | AAG06281.1 | Citronellyl-CoA synthetase | 6e-06 | 21.91 |
| *peg.3130* | Citronellol and citronellal dehydrogenase | Atu | *P. aeruginosa* PAO1 (AtuG) | Citronellol/citronellal dehydrogenase (AtuG) | AAG06280.1 | Citronellal dehydrogenase | 1e-11 | 28.79 |
| *peg.3133* | Enoyl-CoA hydratase | Atu | *P. aeruginosa* PAO1 | Isohexenyl-glutaconyl-CoA hydratase (AtuE) | AAG06278.1 | Isohexenyl-glutaconyl-CoA hydratase | 1e-22 | 30.74 |
| *peg.3137* | 3-Methylcrotonyl-CoA carboxylase alpha subunit | Atu | *P. aeruginosa* PAO1 | Geranyl-CoA carboxylase, α-subunit (AtuF) | AAG06279.1 | Geranyl-CoA carboxylase, α-subunit | 1e-142 | 52.08 |
| *peg.3139* | 3-Methylcrotonyl-CoA carboxylase beta subunit | Atu | *P. aeruginosa* PAO1 | Geranyl-CoA carboxylase, β-subunit (AtuC) | AAG06276.1 | Geranyl-CoA carboxylase, β-subunit | 1e-140 | 43.07 |
| *peg.846* | Hydroxymethylglutaryl-CoA lyase | Liu | *P. aeruginosa* PAO1 | 3-Hydroxy-3-methyl-glutaryl-CoA lyase (LiuE) | AAG05399.1 | 3-Hydroxy-3-methyl-glutaryl-CoA lyase | - | - |
| *peg.847* | Acyl dehydratase | Liu | *P. aeruginosa* PAO1 | 3-Methyl-glutaconyl-CoA hydratase (LiuC) | AAG05401.1 | 3-Methyl-glutaconyl-CoA hydratase | - | - |
| *peg.848* | Methylcrotonoyl-CoA carboxylase subunit alpha | Liu | *P. aeruginosa* PAO1 | Methyl-crotonyl-CoA carboxylase, α-subunit (LiuD) | AAG05400.1 | Methyl-crotonyl-CoA carboxylase, α-subunit | 7e-168 | 59.12 |
| *peg.850* | Methylcrotonoyl-CoA carboxylase beta subunit | Liu | *P. aeruginosa* PAO1 | Methyl-crotonyl-CoA carboxylase, β-subunit (LiuB) | AAG05402.1 | Methyl-crotonyl-CoA carboxylase, β-subunit | 0.0 | 68.54 |
| *peg.852* | Isovaleryl-CoA dehydrogenase | Liu | *P. aeruginosa* PAO1 | Isovaleryl-CoA dehydrogenase (LiuA) | AAG05404.1 | Isovaleryl-CoA dehydrogenase | 2e-175 | 62.43 |
| *peg.853* | Transcriptional regulator | Liu | *P. aeruginosa* PAO1 | Regulator of *Liu* genes (LiuR) | AAG05404.1 | Regulator of *Liu* genes | - | - |

**Table 3.** The most upregulated differentially expressed genes (log_2_FC > 12, -logFDR > 8) in the Lim^+^ cells after 24 h limonene challenge. The complete list of DEGs can be found in Supplementary file S4.

| DEG | Putative function | Contig | logFC | FDR |
| --- | --- | --- | --- | --- |
| *MSTRG.781.1* | no protein identified | NODE_15 | 19.70 | 1.61E-65 |
| *peg.886* | hypothetical protein | NODE_15 | 14.04 | 2.76E-33 |
| *peg.3643* | hypothetical protein | NODE_3 | 13.49 | 5.80E-28 |
| *MSTRG.388.1* | no protein identified | NODE_12 | 13.49 | 3.05E-26 |
| *peg.419* | hypothetical protein | NODE_12 | 13.48 | 4.54E-29 |
| *peg.5044* | pantothenate kinase type III, CoaX-like | NODE_7 | 13.32 | 2.30E-27 |
| *peg.889* | hypothetical protein | NODE_15 | 13.07 | 2.46E-26 |
| *peg.3378* | putative TonB-dependent receptor | NODE_32 | 12.66 | 1.10E-23 |
| *peg.916* | hypothetical protein | NODE_15 | 12.43 | 4.26E-22 |
| *peg.2607* | hypothetical protein | NODE_26 | 12.32 | 2.19E-21 |
| *peg.2393* | MBL-fold metallo-hydrolase superfamily | NODE_23 | 12.32 | 2.66E-21 |
| *peg.915* | transcriptional regulator, TetR family | NODE_15 | 12.30 | 3.27E-21 |
| *peg.985* | hypothetical protein | NODE_15 | 12.30 | 3.46E-21 |
| *peg.3446* | cold shock protein of CSP family | NODE_34 | 12.26 | 4.22E-16 |
| *peg.2714* | hypothetical protein | NODE_27 | 12.23 | 5.10E-21 |
| *peg.2285* | hypothetical protein | NODE_22 | 12.10 | 1.33E-19 |
| *MSTRG.3531.1* | sulfotransferase domain-containing protein | NODE_5 | 12.03 | 8.01E-20 |
| *MSTRG.406.1* | TetR/AcrR family transcriptional regulator | NODE_12 | 12.01 | 1.30E-19 |

**Figure 1.**
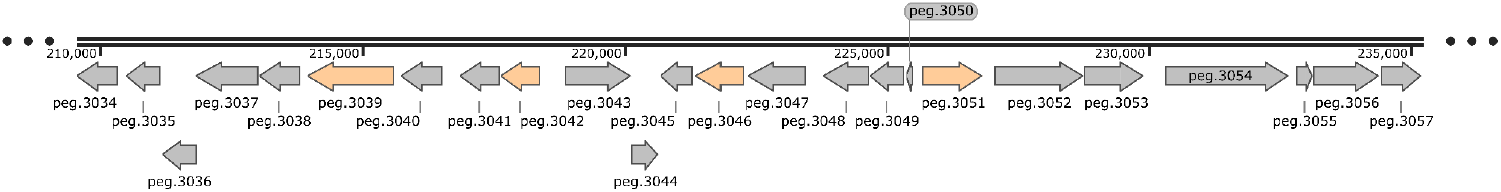
*S. yanoikuyae* A-TP 25 kb genomic cluster (*peg*.*3034* to *peg*.*3057*) arrangement. The putative genes for initial limonene degradation: *peg*.*3039* (1-hydroxy-2-oxo-limonene monooxygenase), *peg*.*3042* (limonene 1,2-diol dehydrogenase) and *peg*.*3051* (limonene monooxygenase) are shown in orange. The full list of the predicted function of the genes in this cluster and their genomic locations can be found in the Supplementary file S2. The figure was created with SnapGene software.

The second cluster, a 30-kb region (*peg*.*3115–peg*.*3144*) (figure 2) contains genes for the acyclic terpene utilization (atu) pathway, responsible for citronellol degradation (5). Unexpectedly, this cluster also harbors genes linked to α-pinene metabolism (*peg*.*3140, peg*.*3124*) (32) despite α-pinene’s structural similarity to limonene rather than citronellol. We also identified genes for leucine and isovalerate degradation (*peg*.*846 - peg*.*853*), by which the product of *atu* pathway is further metabolized to acetyl-CoA (figure S7) (5). To assess whether this genetic repertoire correlates with metabolic activity, *S. yanoikuyae* A-TP was cultivated in liquid medium with citronellol, geraniol, linalool, or β-myrcene as the sole carbon source. Growth was observed only in the presence of citronellol (table S2). The lack of growth on geraniol, and the absence of geranial or geranic acid accumulation in the culture medium suggests that *S. yanoikuyae* A-TP genome does not harbor genes for the initial degradation of this monoterpene. Similarly, the A-TP’s inability to metabolize β-myrcene is likely due to the absence of myrcene hydroxylase, which catalyzes the first reaction in this pathway (6).

**Figure 2.**
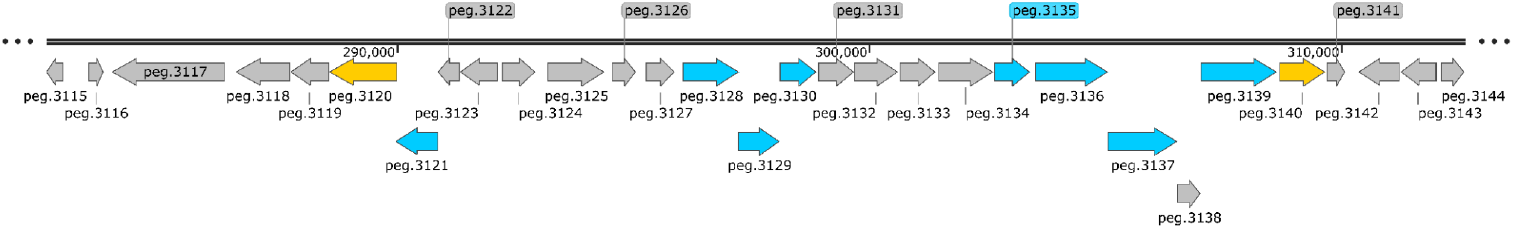
*S. yanoikuyae* A-TP 30 kb genomic cluster (*peg*.*3115* to *peg*.*3144*) arrangement. Putative genes for initial *α*-pinene degradation: *peg*.*3120* (isonovalal dehydrogenase) and *peg*.*3140* (*α*-pinene oxide lyase), and a complete citronellol pathway: *peg*.*3121* (CoA:acetate lyase), *peg*.*3128* (citronellyl-CoA dehydrogenase), *peg*.*3129* (citronellol dehydrogenase), *peg*.*3130* (citronellal dehydrogenase), *peg*.*3135* (isohexenyl-glutaconyl-CoA hydratase), *peg*.*3136* (citronellyl-CoA synthetase), *peg*.*3137* (geranyl-CoA carboxylase, α-subunit), *peg*.*3139* (geranyl-CoA carboxylase, β-subunit), are shown in yellow and blue, respectively. The complete list of the predicted function of the genes in this cluster and their genomic locations can be found in the Supplementary file S2. The figure was created with SnapGene software.

The identification of the two distinct gene clusters, combined with experimental validation of citronellol utilization, provides first insights into the genetic basis of *S. yanoikuyae* A-TP’s monoterpene degradation machinery. This is of special interest as biotechnological production of high-value monoterpene derivatives can contribute to satisfy the growing demand for sustainable and high-quality fragrance ingredients in an economically viable and responsible manner (1).

### Transcriptomic analysis reveals promising candidates for limonene 8-hydratase

In addition to utilization of limonene as a carbon source, *S. yanoikuyae* A-TP also possesses an alternative pathway by which it can biotransform limonene into *α*-terpineol (figure S2). This bioconversion is likely catalyzed by a cofactor-independent limonene 8-hydratase (7). It was reported that the *Geobacillus stearothermophilus* BR388 genome also harbors limonene 8-hydratase (33), however this gene was not found at the reported genomic location (34). A nitrile hydratase from *G. stearothermophilus* was hypothesized to promiscuously catalyze limonene conversion to *α*-terpineol (34), however, no similar proteins were found in *S. yanoikuyae* A-TP.

The lack of publicly available limonene 8-hydratase sequence ruled out the similarity-based approach to identify this enzyme in *S. yanoikuyae* A-TP. However, during the routine maintenance of the bacterium stock on agar plates, we observed the growth of a phenotype that continued to use limonene as a carbon source but lost the ability to convert it to α-terpineol. Therefore, we compared the transcriptional profiles of the two *S. yanoikuyae* A-TP phenotypes, expecting to identify limonene 8-hydratase among differentially expressed genes (figure 3). For clarity, the phenotypes that retained and lost the ability to convert limonene to α-terpineol are hereafter referred to as Lim^+^ and Lim^-^, respectively.

**Figure 3.**
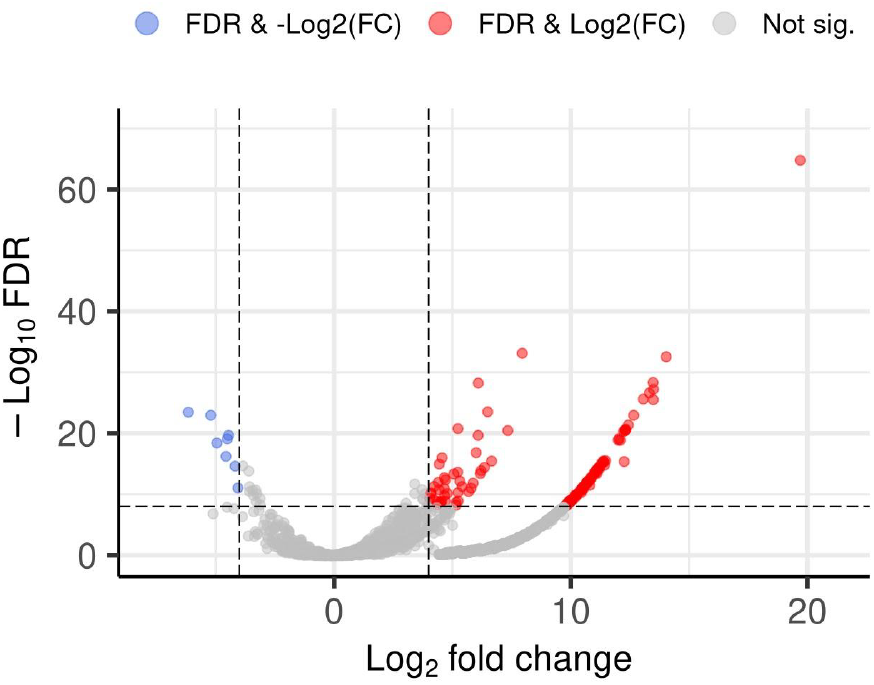
Volcano plot of transcripts expressed in *S. yanoikuyae* A-TP Lim^+^ as compared to Lim^-^ phenotype, after 24 h limonene challenge. Each point on the graph corresponds to one transcript. Upregulated and downregulated transcripts in the Lim^+^ phenotype are shown in red and in blue, respectively, while those in gray were not statistically significant.

The RNA-seq analysis was conducted from triplicate biological samples of both phenotypes, following 24 h incubation with limonene (i.e., limonene biotransformation with resting A-TP cells). An average of 48.1 ± 2.6 and 50.2 ± 2.6 million clean reads was obtained per sample for Lim^+^ and Lim^-^, respectively. The total mapping ratio and uniquely mapped ratio of clean reads were >98.5% and 96%, indicating that sequencing was sufficient to cover all transcripts within the cells. The two phenotypes share >99.4 % 16S rRNA identity, confirming that they are the same species (35).

RNA-seq analysis comparing *S. yanoikuyae* A-TP strains with contrasting α-terpineol production (Lim^+^ vs. Lim^-^) did not identify a clear candidate for limonene 8-hydratase. Among the 95 hydratases (hydro-lyases) encoded by the genome (Supplementary file S3), only *peg*.*5391* (a GDP-mannose 4,6-dehydratase) was upregulated, but its substrate specificity and NADP(H)-dependence make it unlikely to act on limonene.

On the other hand, 1,099 proteins in the A-TP genome lack functional annotation. Considering the lack of publicly available annotation regarding this enzyme, it is likely that limonene 8-hydratase comes from this group of proteins. In fact, three of the ten most upregulated transcripts (*peg*.*3643, peg*.*419*, and *peg*.*916*) have unknown functions and are predicted to contain transmembrane domains (table S3). This makes them promising candidates for future studies as limonene 8-hydratase is suggested to be membrane-associated in *Burkholderia gladioli* (36). *While the most upregulated transcript, mstrg*.*781*, is only 209 pb long and is deemed too short to code for a functional protein, so-called short ORFs (<300 pb), can encode bioactive peptides in bacteria, such as virulence factors and toxins (37). Experimental validation (e.g., gene knockout, heterologous expression) is needed to validate the role of these four candidate genes in limonene biotransformation to *α*-terpineol.

Notably, most monoterpene catabolism genes showed no significant differential expression, which is consistent with the fact that limonene biotransformation to α-terpineol occurs under non-growth conditions (7,8). The two exceptions were *peg*.*3116*, which encodes a short hypothetical protein, and *peg*.*3045*, a TetR-family transcriptional regulator located near limonene degradation genes. In *P. aeruginosa* PAO1, the *atu* operon is regulated by AtuR, a TetR-family repressor that controls citronellol utilization. Therefore, the upregulation of *peg*.*3045* in the Lim^+^ strain may similarly regulate the limonene degradation by redirecting metabolic flux toward α-terpineol production.

## Conclusions

This study provides first insights into the genomic and transcriptomic foundation for the biotechnological use of *Sphingobium yanoikuyae* A-TP. In addition to limonene and α-pinene, the strain utilizes citronellol, with its metabolic genes organized into two distinct clusters. The apparent lack of this strain’s genetic repertoire for geraniol, β-myrcene, and linalool degradation is corroborated by the absence of cell growth on these substrates. On the other hand, we identified four highly upregulated transcripts with unknown functions as strong candidates for limonene 8-hydratase. These results will enable further studies on monoterpene metabolism and support the development of microbial platforms for biotechnological production of monoterpenes and their derivatives.

## Supporting information

Supplementary file S1 - Sphingobium Proteins dataset

Supplementary file S2 - Genes in two genomic clusters

Supplementary file S3 - Hydro-lyases

Supplementary file S4 - Differentially Expressed Genes

## Abbreviations

BVMO: Baeyer-Villiger monooxygenase
DEG: differentially expressed gene
FC: fold change
FDR: False Discovery Rate
GBDP: Genome Blast Distance Phylogeny
GC-MS: gas chromatography-mass spectroscopy
ORF: Open Reading Frame
peg: protein encoding gene
TBDT: Ton-B-dependent transporter
TYGS: Type (Strain) Genome Server.

## CRediT author contribution

N.J. Conceptualization, Formal analysis, Investigation, Methodology, Writing – original draft, Writing – review & editing

G.M.C. Software, Data curation, Methodology, Writing – review & editing

A.L.Q.T. Software, Data curation, Methodology, Writing – review & editing

R.C.M Formal analysis, Investigation, Methodology, Writing – review & editing

M.U.S. Formal analysis, Investigation, Data curation, Writing – review & editing

C.S.C. Methodology, Writing – review & editing

V.L.C. Methodology, Writing – review & editing

J.A.A. Formal analysis, Investigation, Methodology, Writing – review & editing

J.L.B. Supervision, Resources, Writing – review & editing

M.F.R. Supervision, Resources, Project administration, Conceptualization, Writing – review & editing

## Acknowledgements

The authors wish to acknowledge Antonio Jorge da Silva and Daniel Simas for their help with chromatographic analysis, and Claudia Russo and Angela Portella for their help with phylogenetic analysis. Pierre Fontanille is acknowledged for helpful insights and comments on the manuscript.

