## Supplementary Figures and Tables for "Sequencing and functional analysis of *Sphingobium yanoikuyae* A-TP genome reveals genes for utilization of limonene, α-pinene, and citronellol"

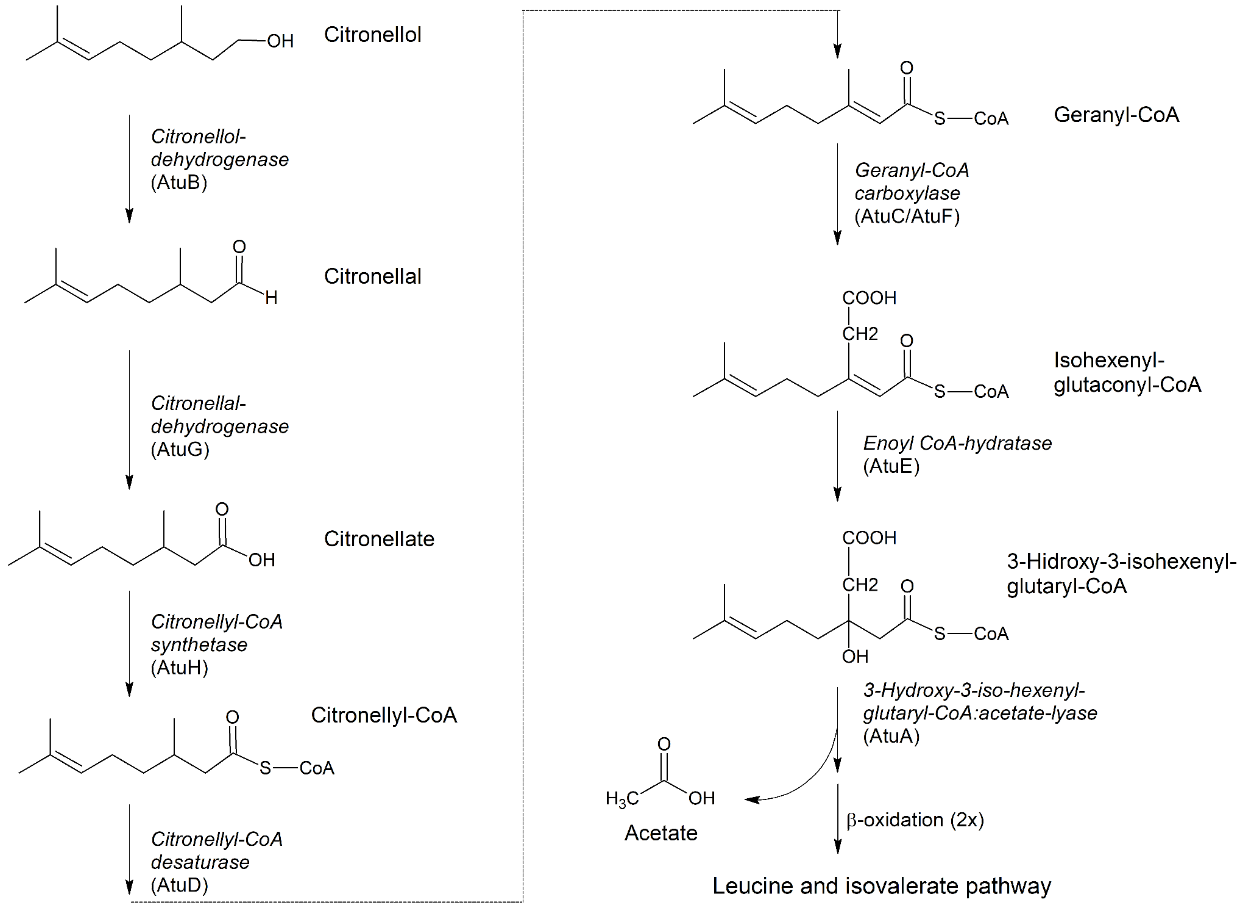


Figure S1. Citronellol pathway and the respective enzymes as described in *Pseudomonas aeruginosa* PAO1 (Adapted from (1)). A degradation begins with the oxidation of citronellol to the corresponding acid (citronelate), which is then converted to a thioester (citronellyl-CoA) and isomerized to geraniol-CoA. A carboxylase transforms the methyl group at the beta position into an acetyl group, giving rise to isohexenyl-glutaconyl-CoA. Next, an enoyl-CoA hydratase eliminates the double bond at the beta position giving rise to 3-hydroxy-3-isohexane glutaryl-CoA. The acetate molecule is then cleaved by an enoyl-CoA lyase resulting in 7-methyl-3-oxo-6-oct enoyl-CoA. This molecule undergoes two rounds of β-oxidation generating methylcrotonyl-CoA, which is sent to leucine and isovaleric acid metabolism.

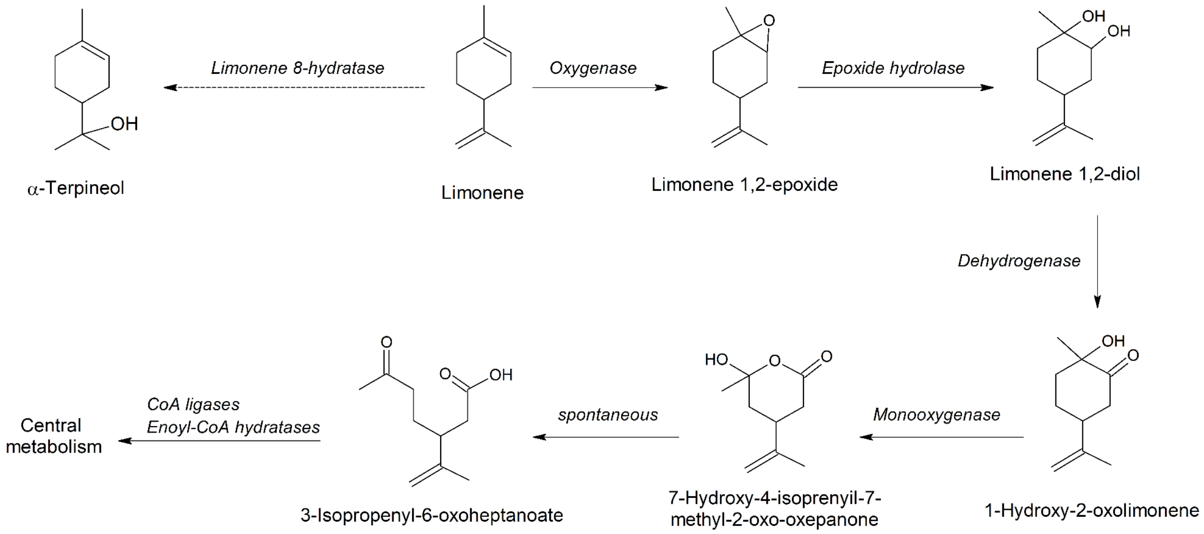


Figure S2. Limonene degradation by *Sphingobium* sp. strain (adapted from (2)). Limonene is first oxygenated to epoxide, which is then hydrolysed to limonene 1,2-diol. A dichlorophenol-indophenol-dependent (DCPIP) dehydrogenase then produces 1-hydroxy-2-oxo limonene, which is then oxygenated by Baeyer-Villiger monooxygenase (BVMO) to yield an unstable lactone that spontaneously rearranges to acyclic 3-isopropenyl-6-oxoheptanoate. This compound is likely further metabolized in reactions analogous to β-oxidation. Limonene conversion to α-terpineol does not produce energy.

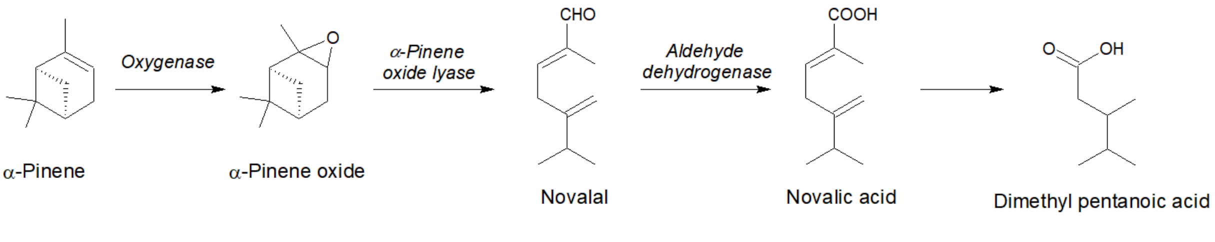


Figure S3. α-Pinene degradation by *Sphingobium* sp. strain (adapted from (2)). The pathway begins with the conversion of α-pinene by a cofactor-dependent monooxygenase. A cofactor-independent lyase (de-cylase) then cleaves both rings of the resulting α-pinene oxide to produce novalal. This molecule is then oxidized by aldehyde dehydrogenase to novalic acid and converted to 3,4-dimethyl pentanoic acid.

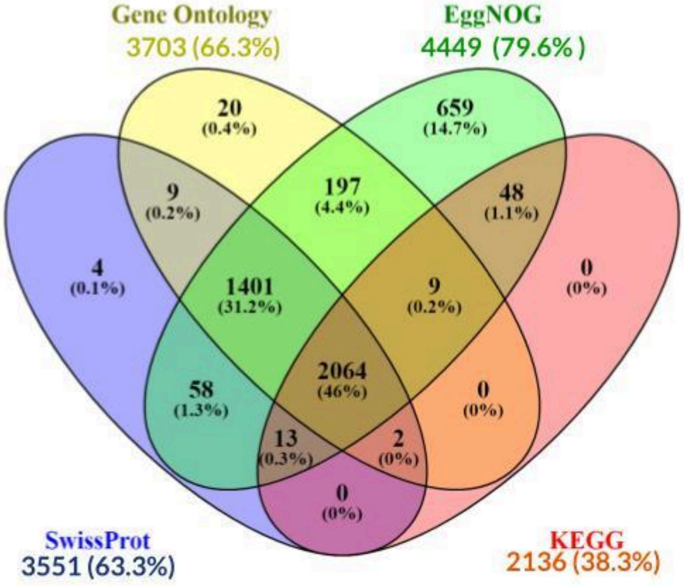


Figure S4. Number of proteins encoded by the *Sphingobium* sp. (*S. yanoikuyae* A-TP) genome in respect to the annotation method. The numbers In parenthesis refer to the percentage of the total predicted proteins.

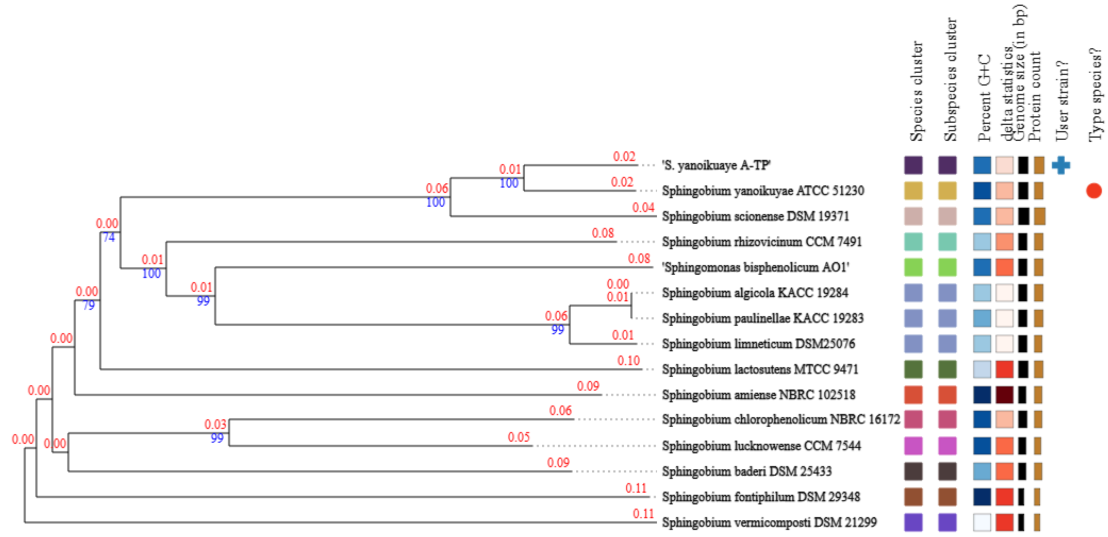


Figure S5. Phylogenomic tree inferred with TYGS web server from GBDP distances calculated from genome sequences (3). Digital DNA–DNA hybridization (dDDH) values and confidence intervals were calculated using the GGDC 3.0 tool's recommended settings (4,5). The resulting intergenomic distances were used to infer a balanced minimum evolution tree with FASTME 2.1.6.1 (6). The branch lengths are scaled in terms of GBDP distance formula $d_5$ (5). The numbers above branches are GBDP pseudo-bootstrap support values > 60 % from 100 replications, with an average branch support of 76.4 %. The tree was rooted at the midpoint (7) and visualized with PhyD3 (8).

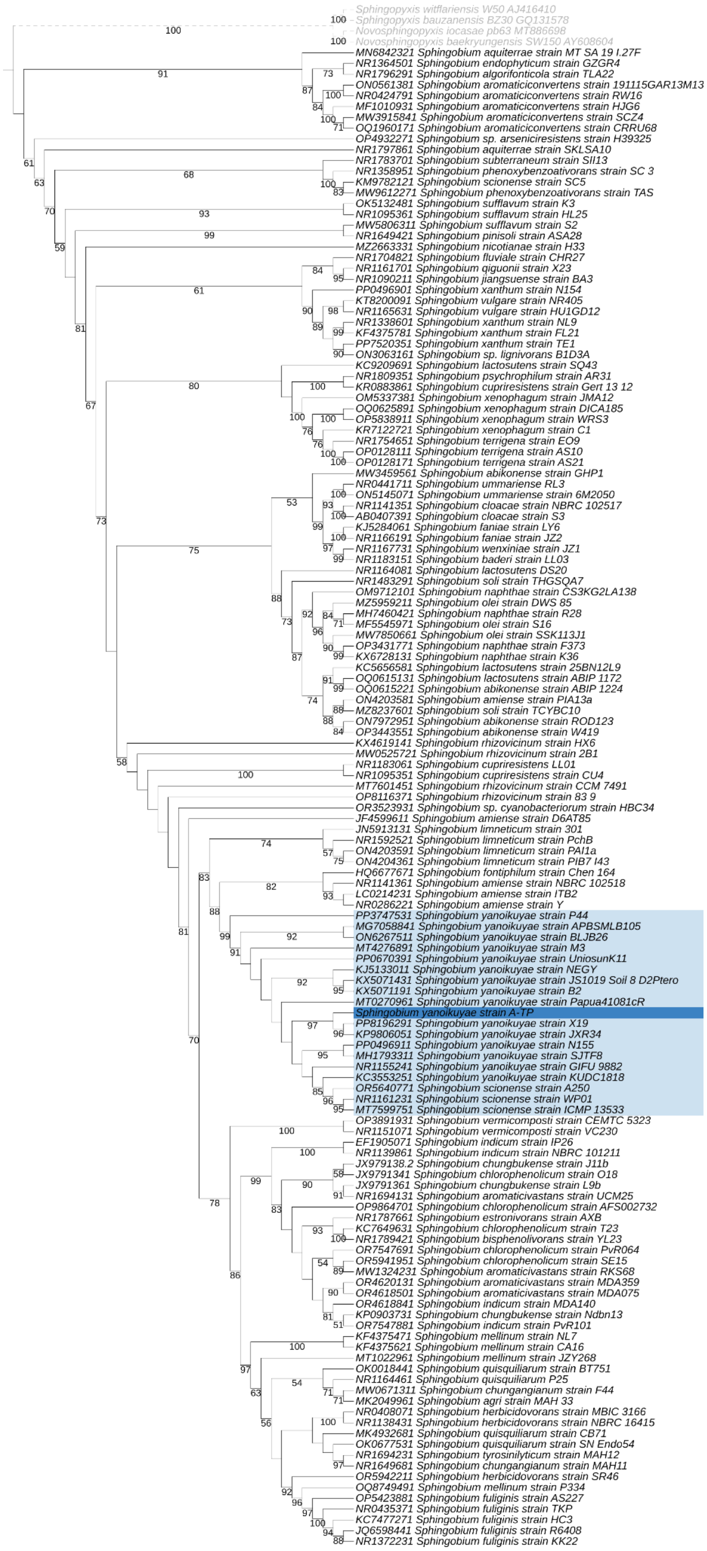


Figure S6. Maximum likelihood phylogenetic tree of *Sphingobium* genus based on the 16S rRNA marker, constructed with 10,000 ultrafast bootstrap replicates. The branches in black indicate bootstrap support values ≥ 50. The light blue clade represents a group composed of sequences from the fifteen *S.*

*yanoikuyae* and three *S. scionense* strains. The sequence of the sample from the present study is highlighted in dark blue.

**Supplementary note:** A phylogenetic tree was created using 16S rRNA sequences from 53 *Sphingobium* species (four strains per species, whenever possible) and from 15 randomly selected *Sphingobium yanoikuyae* strains. The sequences were downloaded from the NCBI database, accessed on 10/06/2024. Sequences from the *Sphingopyxis bauzanensis, Sphingopyxis witflariensis*, *Novosphingopyxis baekryungensis* and *N. iocasae* were used as an outgroup. All sequences were aligned with MAFFT (9), and the alignment was subsequently checked manually to ensure data reliability. The tree was constructed through maximum likelihood analysis using IQ-TREE2 (10), with the evolutionary model selected for optimal fit (K2P+I+G4), and branch support assessed with 10,000 ultrafastbootstrap replicates. The resulting tree was visualized with the iTOL online server (11).

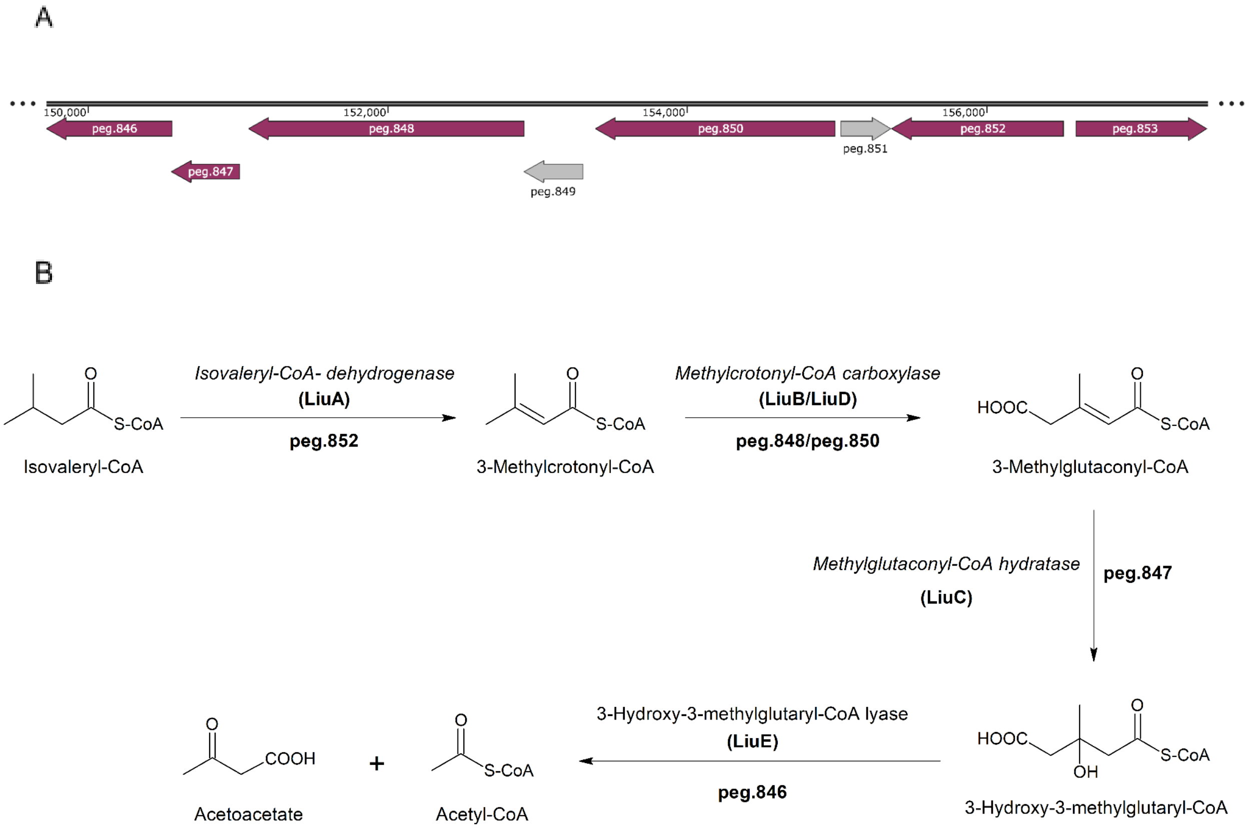


Figure S7. Leucine and isovalerate (Liu) pathway in *S. yanoikuyae* A-TP. (A) Genomic organization of the putative *liu* operon (purple). Function of the genes in gray is unknown. (B) Reactions involved in this pathway are catalyzed by the respective *liu* gene products, as indicated (Adapted from: (1,12))

Table S1. The principal statistical metrics regarding *Sphingobium* sp. *(S. yanoikuyae* A-TP) genome assembly.

| Property | Value | Property | Value |
|---|---|---|---|
| Number of scaffolds | 306 | Number of contigs | 310 |
| Total size of scaffolds | 5669148 bp | Number of contigs in scaffolds | 8 |
| Longest scaffold | 563490 bp | Number of contigs not in scaffolds | 302 |
| Shortest scaffold | 128 bp | Total size of contigs | 5668748 bp |
| Mean scaffold size | 18527 bp | Longest contig | 563490 bp |
| Median scaffold size | 454 bp | Shortest contig | 128 bp |
| N50 scaffold length | 199910 bp | N50 contig length | 199910 bp |
| L50 scaffold count | 8 | L50 contig count | 8 |

Table S2. Growth of *S. yanoikuyae* A-TP cells in the presence of different acyclic monoterpenes.

| **Monoterpene** | **Pre-cultures with 2 % (v/v) monoterpene** | **Cultures with 1 % (v/v) monoterpene** |
|---|---|---|
| Citronellol | + | + |
| Geraniol | - | - |
| Myrcene | - | - |
| Linalool | - | - |

Table S3. Differentially expressed transcripts (DETs) in $Lim^+$ phenotype with predicted transmembrane domains. Symbol (*) corresponds to the sole DET with predicted signal peptide.

| ID | Putative annotation | *Contig* | logFC | FDR |
|---|---|---|---|---|
| *peg.3643* | Hypothetical protein | NODE_3 | 13.49 | 5.80E-28 |
| *peg.419* | Hypothetical protein | NODE_12 | 13.48 | 4.54E-29 |
| *peg.3378* | Putative TonB-dependent receptor | NODE_32 | 12.66 | 1.10E-23 |
| *peg.916* | Hypothetical protein | NODE_15 | 12.43 | 4.26E-22 |
| *peg.2607* | Hypothetical protein | NODE_26 | 12.32 | 2.19E-21 |
| *MSTRG.3531.1* | Sulfotransferase domain-containing protein | NODE_5 | 12.03 | 8.01E-20 |
| *MSTRG.906.1* | Major facilitator superfamily (MFS) transporter | NODE_16 | 11.20 | 4.00E-15 |
| *peg.5119* | Hypothetical protein | NODE_7 | 11.12 | 1.67E-14 |
| *peg.1961* | Glycosyltransferase, group 2 family protein | NODE_1 | 10.98 | 3.74E-14 |
| *peg.332* | Hypothetical protein | NODE_11 | 10.86 | 1.61E-13 |
| *peg.2179* | Hypothetical protein | NODE_21 | 10.81 | 3.07E-12 |
| *peg.873* | Quinate/shikimate dehydrogenase | NODE_15 | 10.69 | 1.20E-12 |
| *MSTRG.1446.1** | Histidinol-phosphate/aromatic aminotransferase | NODE_1 | 10.68 | 1.03E-12 |
| *peg.2295* | Hypothetical protein | NODE_22 | 10.30 | 7.43E-11 |
| *peg.440* | HlyD-like membrane fusion protein YhiI | NODE_12 | 9.97 | 1.23E-09 |
| *MSTRG.4233.1* | Hypothetical protein | NODE_8 | 7.95 | 7.24E-34 |
| *peg.3251* | Fatty acid desaturase, type 2 | NODE_2 | 5.32 | 6.55E-13 |
| *peg.1144* | Hypothetical protein | NODE_16 | 5.24 | 5.08E-11 |
| *peg.3519* | Hypothetical protein | NODE_3 | 4.66 | 1.81E-13 |
| *peg.1568* | Glucans biosynthesis glucosyltransferase H | NODE_1 | 4.56 | 1.38E-09 |
| *peg.91* | MotA/TolQ/ExbB proton channel family protein | NODE_10 | -4.46 | 2.13E-20 |
| *peg.90* | Biopolymer transport protein ExbD/TolR | NODE_10 | -4.51 | 8.01E-20 |
